# RNA chaperones Hfq and ProQ play a key role in the virulence of the plant pathogenic bacterium *Dickeya dadantii*

**DOI:** 10.1101/2021.03.26.437193

**Authors:** Simon Leonard, Camille Villard, William Nasser, Sylvie Reverchon, Florence Hommais

**Affiliations:** Université de Lyon, INSA-Lyon, Université Claude Bernard Lyon1, CNRS, UMR5240 MAP, Microbiologie, Adaptation, Pathogénie, F-69622, Villeurbanne CEDEX, France

**Keywords:** *Dickeya dadantii*, *proQ*, *hfq*, virulence, non coding RNA

## Abstract

*Dickeya dadantii* is an important pathogenic bacterium that infects a number of crops including potato and chicory. While extensive works have been carried out on the control of the transcription of its genes encoding the main virulence functions, little information is available on the post-transcriptional regulation of these functions. We investigated the involvement of the RNA chaperones Hfq and ProQ in the production of the main *D. dadantii* virulence functions. Phenotypic assays on the *hfq* and *proQ* mutants showed that inactivation of *hfq* resulted in a growth defect, a modified capacity for biofilm formation and strongly reduced motility, and in the production of degradative extracellular enzymes (proteases, cellulase and pectate lyases). Accordingly, the *hfq* mutant failed to cause soft rot on chicory leaves. The *proQ* mutant had reduced resistance to osmotic stress, reduced extracellular pectate lyase activity compared to the wild-type strain, and reduced virulence on chicory leaves. Most of the phenotypes of the *hfq* and *proQ* mutants were related to the low amounts of mRNA of the corresponding virulence factors. Complementation of the double mutant *hfq-proQ* by each individual protein and cross-complementation of each chaperone suggested that they might exert their effects via partially overlapping but different sets of targets. Overall, it clearly appeared that the two Hfq and ProQ RNA chaperones are important regulators of pathogenicity in *D. dadantii*. This underscores that virulence genes are regulated post transcriptionally by non-coding RNAs.

## 1 Introduction

*Dickeya dadantii* is a Gram-negative phytopathogenic bacterium responsible for soft rot disease in a wide range of plant species including economically important crops (e.g. potato, chicory, sugar beet) and many ornamental plants (Ma et al., 2007). It causes important production losses (Toth et al., 2011). Virulence mechanisms of *D. dadantii* have been extensively studied (Ma et al., 2007). The infection process is divided in two distinct phases: (i) an asymptomatic phase when the bacterium penetrates into the host and progresses through intercellular spaces without multiplying substantially; (ii) a symptomatic phase associated with strongly increased bacterial fitness and multiplication (Fagard et al., 2007). Globally, the four main steps of plant infection by *Dickeya* are the following: (i) adherence to the plant surface and entry into the plant tissues, via wound sites or through natural openings such as stomata, (ii) colonization of the apoplastic spaces between plant cells, (iii) suppression of the host defense response, and (iv) plant cell wall degradation (through degradative extracellular enzyme production, mainly pectate lyases) resulting in the development of disease symptoms. Each of these disease stages and life-history transitions requires the correct spatio-temporal production of the different adaptive and virulence factors (including those involved in adhesion, motility, stress resistance and plant cell wall degradation) in response to various signals (changes in cell density, variation in environmental physico-chemical parameters, and host disease reaction) (Reverchon and Nasser, 2013).

To characterize the regulation of this pathogenic process, investigations on *D. dadantii* have mostly focused on its control by DNA-binding transcription factors (Reverchon and Nasser, 2013; Leonard et al., 2017) with a few additional studies about the regulatory role of chromosome dynamics (Ouafa et al., 2012; Jiang et al., 2015; Meyer et al., 2018). Knowledge of the post-transcriptional regulation of virulence factor production by sRNAs in *D. dadantii* is still in its infancy.

Post-transcriptional regulation is defined as the control of gene expression at the RNA level and classically occurs through base-pairing interactions between regulatory RNAs (sRNAs) and mRNAs. This base pairing can have positive or negative effects on the stability and/or the translation of the targeted mRNA. These sRNAs can be broadly divided into two categories according to their genomic location: (i) *cis*-acting antisense sRNAs are transcribed from the opposite strand of their targets and act via extensive base pairing; (ii) *trans*-acting sRNAs mostly originate from intergenic regions, display partial sequence complementarities with their mRNA targets and can regulate more than one target. The interactions between sRNAs and their targets are often assisted by specialized RNA-binding proteins called RNA chaperones.

A prominent bacterial RNA chaperone is the Hfq protein which contributes to regulation by *trans*-acting sRNAs in many bacteria (Updegrove et al., 2016). Hfq was first discovered in *Escherichia coli* as an essential host factor of the RNA bacteriophage Qbeta. Hfq impacts multiple steps, like changing RNA structure, bringing RNAs into proximity, neutralizing the negative charge of the two pairing RNAs, stimulating the nucleation of the first base pairs as well as facilitating the further annealing of the two RNA strands. While estimates of the number of Hfq vary from ≈20,000 to 60,000 (Kajitani et al., 1994; Ali Azam et al., 1999), it is clear that Hfq is limiting under most conditions (Wagner, 2013).

Other proteins with possible chaperone activity have been reported recently. For example, the monomeric ProQ protein of *Salmonella enterica* is an RNA-binding protein that interacts with and stabilizes over 50 highly structured antisense and *trans*-acting sRNAs. (Smirnov et al., 2016). The cellular concentration of ProQ was estimated to be 2,000 copies per cell (Sheidy and Zielke, 2013). This protein was originally identified as being important for osmolyte accumulation in *E. coli* by increasing cellular levels of the proline transporter ProP (Milner and Wood, 1989; Kunte et al., 1999) and was later shown to possess RNA strand exchange and RNA annealing activities (Chaulk et al., 2011). Thus, ProQ was initially described as an RNA chaperone that controls ProP levels in *E. coli*. In *Legionella pneumophilia*, the ProQ equivalent protein (called RocC) interacts with one *trans*-acting sRNA to control the expression of genes involved in natural transformation (Attaiech et al., 2016).

ProQ belongs to the RNA-binding proteins of the FinO family. FinO has been studied for its role as an RNA chaperone in antisense regulation of F plasmid conjugation in *E. coli* (Mark Glover et al., 2015). As shown in *S. enterica*, ProQ seems to recognize stable RNA hairpins such as transcriptional terminators and reading the RNA structure rather than its sequence (Holmqvist et al., 2018).

While several recent studies have addressed a potential role of Hfq in the virulence of phytopathogenic bacteria like *Agrobacterium tumefaciens* (Wilms et al., 2012), *Erwinia amylovora* (Zeng et al., 2013), *Pectobacterium carotovorum* (Wang et al., 2018) and *Xanthomonas campestris* (Lai et al., 2018), nothing is known about the impact of Hfq and ProQ on *D. dadantii* virulence. Moreover, potential links between ProQ and the virulence of plant-pathogenic bacteria have never been established. To address these questions, we constructed and characterized *hfq* and *proQ* mutants. Loss of Hfq or ProQ resulted in drastically reduced virulence. This phenotype was associated with the alteration of several virulence determinants including pectate lyase production, motility, and adhesion. Additionally, analyses of mutants defective in the two proteins suggested that these two RNA chaperones might exert their effects via partially overlapping but different sets of targets.

## 2 Materials and Methods

### 2.1 Bacterial strains, plasmids and culture conditions

The bacterial strains, plasmids, phages and primers used in this study are described in Tables S1, S2 and S3. *E. coli* and *D. dadantii* strains were grown at 37°C and 30°C, respectively, in Luria-Bertani broth (LB) medium or in M63 minimal medium (Miller, 1972) supplemented with 0.1 mM CaCl_2_, 0.2% (w/v) sucrose and 0.25% (w/v) polygalacturonate (PGA, a pectin derivative) as carbon sources. PGA induces the synthesis of pectate lyases, which are the essential virulence factors of *D. dadantii*. When required, the media were supplemented with antibiotics at the following concentrations: ampicillin (Amp) 100 μg/mL, chloramphenicol (Cm) 20 μg/mL, kanamycin (Kan) 50 μg/mL. The media were solidified with 1.5 % (w/v) Difco agar. Liquid cultures were grown in a shaking incubator (220 r.p.m.). Bacterial growth in liquid medium was estimated by measuring turbidity at 600 nm (OD_600_) to determine growth rates.

### 2.2 Gene knockout and complementation of the Hfq- and ProQ-encoding genes in *D. dadantii*

The *hfq* gene was inactivated by introducing a *uidA-Kan* cassette into the unique *BsrG*I restriction site present in its open reading frame. The *uidA-Kan* cassettes (Bardonnet and Blanco, 1992) includes a promoterless *uidA* gene that conserves its Shine Dalgarno sequence.

To create a Δ*proQ::Cm* mutant, segments located 500 bp upstream and 500 bp downstream of *proQ* were amplified by PCR using primer pairs P1-P2 and P3-P4 (Table S3). Primers P2 and P3 included a unique restriction site for *BglII* and were designed to have a short 20-bp overlap of complementary sequences. The two separate PCR fragments were attached together by overlap extension polymerase chain reaction using primers P1 and P4. The resulting Δ*proQ-BglII* PCR product was cloned into a pGEMT plasmid to create plasmid pGEM-T-ΔproQ-BglII. The Cm resistance cassette from plasmid pKD3 (Datsenko and Wanner, 2000) was inserted into the unique *BglII* site of pGEM-T-ΔproQ-BglII to generate pGEM-T-ΔproQ::Cm (Table S2).

We took care to select cassettes without transcription termination signals in order to avoid polar effects on downstream genes for both insertions. The insertions were introduced into the *D. dadantii* chromosome by marker exchange recombination between the chromosomal allele and the plasmid-borne mutated allele. The recombinants were selected after successive cultures in low phosphate medium in the presence of the suitable antibiotic because pBR322 derivatives are very unstable in these conditions (Roeder and Collmer, 1985). Correct recombination was confirmed by PCR. Mutations were transduced into a clean *D. dadantii* 3937 genetic background using phage ΦEC2 (Table S1).

For complementation of the *hfq* and *proQ* mutations, the promoter and coding sequences of the *proQ* and *hfq* genes were amplified from *D. dadantii* 3937 genomic DNA using primers P5/P6 and P7/P8, respectively (Table S3). The forward primers (P5 and P7) included a unique restriction site for *NheI*, and the reverse primers (P6 and P8) included a unique restriction site for *HindIII*. After digestion with *NheI* and *HindIII*, each PCR fragment was ligated into pBBR1-mcs4 previously digested by *NheI* and *HindIII* to generate *pBBR1-mcs4::hfq* and *pBBR1-mcs4::proQ*, respectively (Table S2). Correct constructions were confirmed by sequencing.

### 2.3 Agar plate detection tests for pectate lyase, cellulase, protease and other enzyme assays

Protease activity was detected on medium containing skim milk (12.5 g L^-1^). Cellulase activity was detected on carboxymethylcellulose agar plates with the Congo red staining procedure (Teather and Wood, 1982). Pectate lyases were assayed on toluenized cell extracts. Pectate lyase activity was measured by recording the degradation of PGA into unsaturated products that absorb at 230 nm (Moran et al., 1968). Specific activity was expressed as nmol of unsaturated products liberated per min per mg of bacterial dry weight, given that an OD_600_ of 1 corresponded to 10^9^ bacteria.mL^-1^ and to 0.47 mg of bacterial dry weight.mL^-1^.

### 2.4 Stress resistance assays

Bacteria were cultured at 30°C in 96-well plates using M63S (M63 + 0.2% w/v sucrose), pH 7.0, as minimal medium. Bacterial growth (OD_600_) was monitored for 48 h using an <u>Infinite^®^ 200 PRO - Tecan</u> instrument. Resistance to osmotic stress was analyzed using M63S enriched in 0.05 to 0.5 M NaCl. Resistance to oxidative stress was analyzed in the same medium by adding H_2_O_2_ concentrations ranging from 25 to 200 μM. The pH effect was analyzed using the same M63S medium buffered with malic acid at different pH values ranging from 3.7 to 7.0.

### 2.5 Biofilm measurements

Biofilm formation was quantified using the microtiter plate static biofilm model. Bacteria were grown for 48 h at 30°C in 24-well plates in M63 medium supplemented with glycerol as the carbon source. Then, the supernatant was removed, and the biofilm was washed once with 1mL of M63 medium and resuspended in 1 mL of the same medium. The percentage of adherence was then calculated as the ratio of the number of cells in the biofilm over the total number of cells, i.e. biofilm cells over planktonic cells. The amount of planktonic cells was estimated by measuring the optical density at 600 nm of the supernatant and the washing buffer. The amount of cells in the biofilm was estimated by measuring the OD_600_ of the biofilm resuspended in M63.

### 2.6 Motility and chemotaxis assays

For the *proQ* mutant, motility assays were performed on semi-solid LB agar plates. An overnight bacterial culture was prepared as described above, and then inoculated in the centre of each of eight Petri dishes with a sterile toothpick. For motility experiments, 0.3% agar plates were used. Halo sizes were examined after incubation at 30°C for 24 h. Motility indexes were calculated as the ratios of the mutant halo size over the wild type (WT) halo size.

For the *hfq* mutant, motility assays were performed in competition (to avoid the influence of bacterial growth), as previously described (Ashby et al., 1988). Briefly, 10 mL of bacteria in their exponential growth phase were washed twice in sodium-free buffer and then concentrated in 3 mL. Capillary assays were performed in competition in an equal 1:1 ratio. Suspension dilutions of chemotaxis assays were spotted onto selective LB agar medium. Different bacterial populations were thus enumerated on LB agar plates (both wild-type cells and *hfq::uidA*-Kan mutants) and LB agar plates containing kanamycin (*hfq::uidA*-Kan mutants). Motility indexes were calculated as the ratios of the number of *hfq::uidA*-Kan mutants over the number of wild-type cells.

### 2.7 Virulence assays

Virulence assays were performed on wounded chicory leaves by depositing a drop of inoculum as previously described (Dellagi et al., 2005). Briefly, chicory leaves were wounded with a 2 cm incision using a sterile scalpel, inoculated with 5 μL of bacterial suspension (OD_600_=1) and incubated at 30°C in a dew chamber at 100% relative humidity. Disease severity was determined 18 h and 48 h after inoculation by collecting and weighing the macerated tissues.

### 2.8 Quantitative RT-PCR analyses

Gene expression analyses were performed using qRT-PCR. Total RNAs were extracted and purified from cultures grown to the late exponential phase (OD_600_ = 0.8) as previously described (Maes and Messens, 1992; Hommais et al., 2008). Reverse transcription and quantitative PCR were performed using the REvertAid First Strand cDNA synthesis kit and the Maxima SYBR Green/ROX qPCR Master Mix (Thermo Scientific) with an LC480 Lightcycler (Roche). Primer specificity was verified by melting curve analysis. qPCR primers are listed in Table S4.

### 2.9 Data representation and statistical analysis

Boxplot representations were generated using R software (R Core Team, 2020) and the beeswarm package (Eklund, 2016). Statistical analysis was performed using Wilcoxon Mann-Whitney tests, and differences were considered significant when the *p* value < 0.05.

## 3 Results

### 3.1 Analysis of *D. dadantii* Hfq and ProQ protein sequences and their genomic contexts

*E. coli* and *D. dadantii* Hfq proteins displayed 83% identity. The highest identity level was in the N-terminal region (amino acids 1-74), which forms the core of the protein and contains its RNA-binding sites (Link et al., 2009). Most of the amino acids involved in RNA interactions were conserved except E18, which was K18 in *D. dadantii* (Figure S1). *D. dadantii* ProQ was 68% identical with *E. coli* ProQ, with also high identity in the N-terminal FinO domain of ProQ, which is the primary determinant of its RNA-binding capacity (Chaulk et al., 2011; Gonzalez et al., 2017). In particular, the regions spanning residues 1-10 and 92-105, shown to interact with RNA, were highly conserved (Gonzalez et al., 2017). All the residues involved in the formation of a basic patch on the protein surface (R32, R69, R80, R100, K101, K107, and R114) – an important structure for interaction with RNAs – were conserved (Figure S1).

The *hfq* and *proQ* genes are embedded in the same chromosomal context in *D. dadantii* as in other bacteria such as *E. coli* or *S. enterica* (Figure S2). The *hfq* gene is part of the well conserved *amiB-mutL-miaA-hfq-hflXKC* cluster (Tsui and Winkler, 1994), while *proQ* is localized between *yebR* and *prc*. Inspection of the transcriptomes of *D. dadantii* under various physiological conditions (Reverchon et al., 2021) showed that transcription of *hfq* could be driven by (i) a promoter upstream of *mutL*, (ii) a promoter inside *mutL* and upstream of *miaA*, or (iii) two promoters inside *miaA* and upstream of *hfq* (Figure S2). Considering the expression level of *mutL-miaA-hfq* genes, it appears that *hfq* was largely transcribed from the two promoters inside *miaA* and rarely co-transcribed with *miaA*. The downstream genes showed similar expression profiles and did not exhibit any promoter activity downstream of *hfq*, suggesting that they may be co-transcribed with *hfq* in the same way as in *E. coli* (RegulonDB, http://regulondb.ccg.unam.mx/). Two promoters were found upstream of the *proQ* gene (one between*proQ* and *yebR* and one upstream of *yebR)* (Figure S2B). Regarding the difference in read coverage obtained from RNA-seq experiments, *yebR* and *proQ* seemed to be largely transcribed separately (Figure S2B). On the contrary, *prc* and *proQ* had similar coverage, and no transcription start site was found between them, supporting co-transcription similarly to what is observed in *E. coli* (RegulonDB, http://regulondb.ccg.unam.mx/).

### 3.2 Phenotypic characterization of the *hfq* and *proQ* mutants

We first analyzed the growth characteristics of the *hfq* and *proQ* mutants. The WT, *hfq* and *proQ* strains were grown in LB rich medium and in M63 minimal medium supplemented with sucrose as the sole carbon source. While the *proQ* mutant and the WT grew similarly in both media, the *hfq* mutant exhibited delayed growth. However, in rich medium both the WT strain and *hfq* mutant reached the same optical density after being grown for 26 hrs (Figure 1A). In minimal medium with sucrose as the sole carbon source, the *hfq* mutant grew much more slowly than the WT, and reached the stationary phase at a lower optical density (Figure 1B). The growth defect of the *hfq* mutant was fully restored by complementation with plasmid pBBR-mcs4:: *hfq* (Figure 1), indicating that the *hfq::uidA*-Kan insertion had no polar effects on downstream *hflXKC* genes. In contrast, transformation of the *proQ* mutant and WT strains with the pBBR-mcs4::*proQ* plasmid expressing *proQ* led to a lower growth rate, especially in minimal medium (Figure 1D). The two strains grew similarly in the absence of the pBBR-mcs4::*proQ* plasmid. This suggests that slight ProQ overexpression compromises growth irrespective of the genetic background. These data demonstrate that *hfq* mutation retards cellular growth, while *proQ* mutation does not. The effect was more pronounced in minimal medium compared to rich medium, suggesting that Hfq plays a more important role in the ability of *D. dadantii* to grow under conditions of nutrient limitation. A similar growth defect of *hfq* mutants has been observed in other bacteria such as *P.carotovorum* (Wang et al., 2018), *A. tumefaciens* (Wilms et al., 2012) *or E. amylovora* (Zeng et al., 2013).

**Figure 1:**
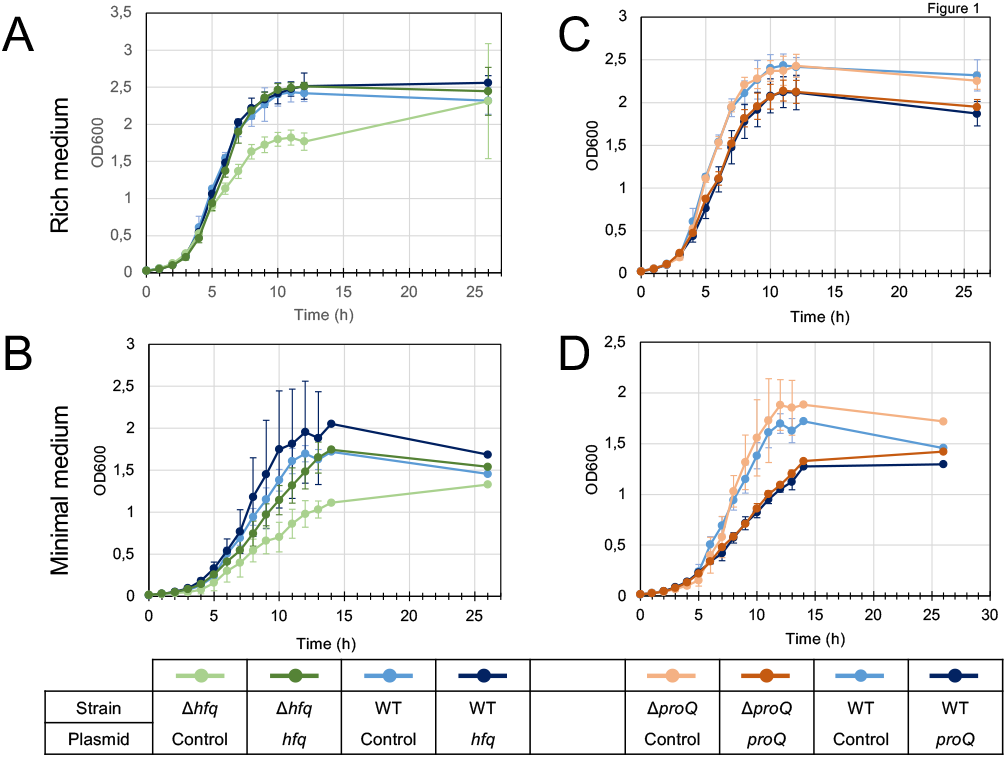
Growth of the wild type, mutant and complemented strains in LB rich medium (A and C) and M63 minimal medium supplemented with sucrose (B and D). Overnight bacterial precultures were diluted to an OD_600_ of 0.03 in the same growth medium. OD_600_ measurements of the culture were made at regular intervals to determine growth rates. The experiment was repeated three times. The graph shows curves from one representative experiment.

*Dickeya* encounter various stresses during their pathogenic growth, so we assessed the stress resistance of the *hfq* and *proQ* mutants (Figure S3). They both showed behaviors similar to the WT strain regarding pH and H_2_O_2_ stress resistance, but displayed a higher sensitivity to osmotic stress than the WT strain did. The *hfq* mutant displayed a 50% growth rate reduction on 0.4 M NaCl, while the WT strain was only slightly affected (20% growth rate reduction). This effect was even more pronounced at 0.5 M NaCl, with a growth rate reduction of 90 % for *hfq* compared to 60% for the WT. The *proQ* mutant did not grow at 0.3M NaCl and at higher NaCl concentrations (Figure S3). Complementation experiments revealed that expression of *hfq* or *proQ* from an episome (plasmid pBBR-mcs4::*hfq* and pBBR1-mcs4::*proQ*) fully restored the osmotic resistance of the two mutants to the WT level (Figure S3). We inferred that the two chaperones are involved in providing resistance to osmotic stress. Overall, this result is consistent but not identical with previous studies showing that Hfq and ProQ contribute to stress tolerance, including nutrient deprivation, osmotic stress and oxidative stress in *Salmonella* and *E. coli* (Chaulk et al., 2011; Smirnov et al., 2017).

### 3.3 Hfq and ProQ are required for full virulence of *Dickeya dadantii*

The virulence of the *hfq* and *proQ* mutants was tested on chicory leaves. The *hfq* mutant was severely impaired in virulence, and soft rot symptoms were drastically reduced (Figure 2A). Disease symptoms were observed following inoculation with the *proQ* mutant and the WT strain, but they were less severe in the *proQ* background. Quantitative results obtained by measuring the weight of macerated tissues showed a significant difference between the disease symptoms induced by each strain (p-value = 1.5e-3) (Figure 2B & C). As observed previously, the virulence defect of the *hfq* mutant was more severe than that of the *proQ* mutant, in as far as the *hfq* mutant did not exhibit any macerated tissue (p-value = 5.7e-4). Consequently, we did not weigh any macerated tissue in the *hfq* mutant. The soft rot symptoms caused by the *hfq* and *proQ* mutants did not increase over longer incubation times (48 h). The lower virulence of *hfq* and *proQ* was therefore not solely related to retarded cellular growth. Complementation experiments revealed that expression of *hfq* from an episome (plasmid pBBR-mcs4::*hfq*) fully restored the impaired virulence of the *hfq* mutant (Figure 2B). Likewise, expression of *proQ* from an episome (plasmid pBBR-mcs4::*proQ*) restored the impaired virulence of the *proQ* mutant (Figure 2C).

**Figure 2:**
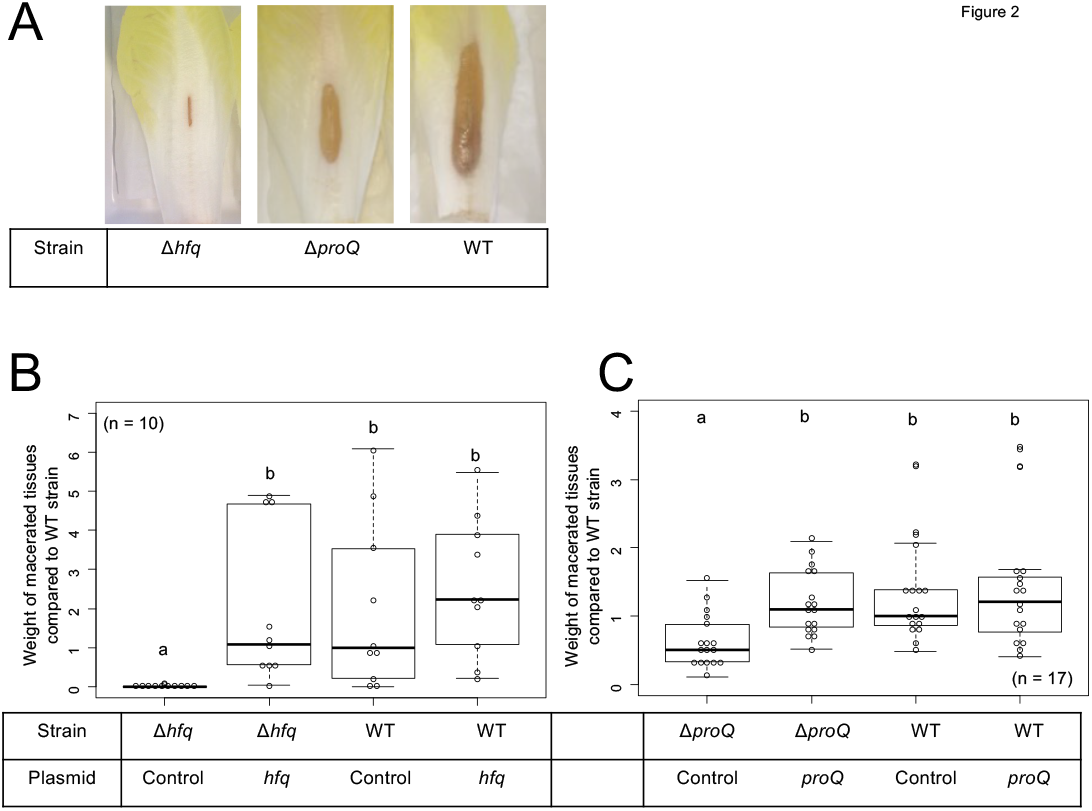
Impact of Hfq and ProQ on *D. dadantii* virulence. A, Representative examples of symptoms induced by the wild type and mutant strains; B and C, weights of macerated tissues following infection by the *hfq* and *proQ* mutants, the wild type strain and the complemented strains. Data were normalized based on the weights of macerated tissues from the wild-type strain. Chicory leaf assays were performed as described in the Materials and methods section with an incubation time of 18h, and weights of macerated tissues were measured. a/b/c/d boxplot annotations highlight significant differences (*P* < 0.05, Wilcoxon Mann-Whitney test).

Thus, the *hfq* and *proQ* genes are required for efficient pathogenic growth since both mutants were clearly impaired in initiating maceration and inducing soft rot symptoms, albeit to different extents.

### 3.4 Production of late virulence factors, pectate lyase, protease and cellulase, is abolished in the *hfq* mutant and reduced in the *proQ* mutant

*D. dadantii* is known to use several essential virulence factors that collectively contribute to its ability to cause disease. These factors include production of cell-wall-degrading enzymes like pectate lyases, proteases and cellulase, which are responsible for soft rot symptoms. To clarify whether Hfq and ProQ have any influence on the production of key virulence factors, we compared enzyme activity in *hfq* and *proQ* mutant extracts with WT strain extracts (Figure 3A). Pectate lyase activity was abolished in the *hfq* mutant (*p*-value = 2.4e-7). This defect in pectate lyase activity was not a consequence of the growth defect of the *hfq* mutant since activities were normalized to cell density. Also, the levels of pectate lyase activity were significantly reduced in the *proQ* mutant compared to the WT (*p*-value = 2.0e-9) (Figure 3A). Reduced cellulase and protease activities were also observed on carboxymethylcellulose and skim milk agar plates for each mutant (Figure 3B). Complementation experiments showed that the impaired production of cell-wall-degrading enzymes was restored in *hfq* and *proQ* mutant cells transformed with the pBBR-mcs4::*hfq* and pBBR-mcs4::*proQ* plasmids, respectively (Figure 3). Therefore, the impaired pathogenicity of *D. dadantii* due to *hfq* and *proQ* mutations is linked with reduced production of late virulence factors in these mutant strains.

**Figure 3:**
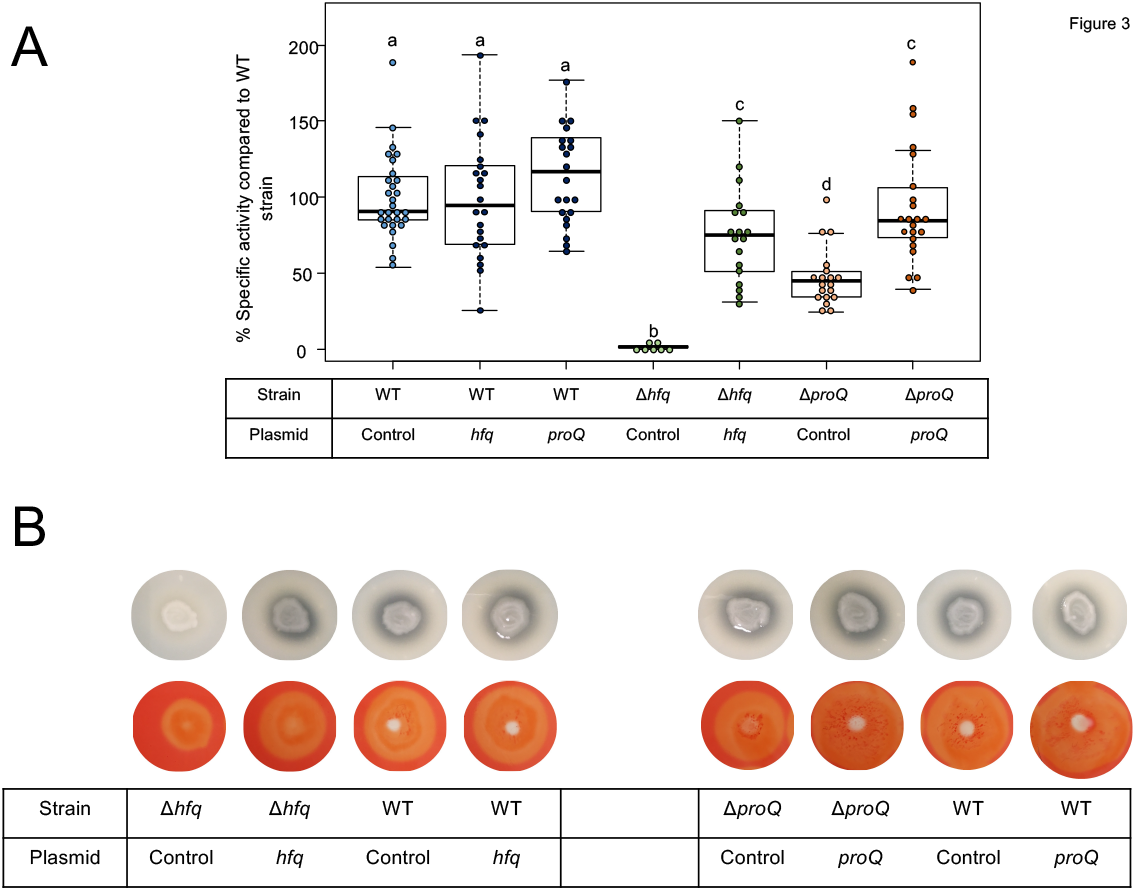
Impact of Hfq and ProQ on cell-wall-degrading enzymes. A, Pectate lyase activity in the wild type, mutant and complemented strains, expressed as percent of wild type strain specific activity. a/b boxplot annotations highlight significant differences (*P* < 0.05, Wilcoxon Mann-Whitney test). B, protease production on medium containing skim milk, and cellulase production on carboxymethylcellulose agar plate with Congo red staining in the *hfq* and *proQ* mutants, the complemented strains and the wild type strain.

### 3.5 Early virulence determinants such as biofilm production and motility are also impaired in the *hfq* and *proQ* mutants

At the initial stage of infection, *D. dadantii* must adhere to the plant surface and enter into the apoplast. *D. dadantii* produces cellulose fibrils, which enable it to develop aggregates on the plant surface (Jahn et al., 2011; Prigent-Combaret et al., 2012). These aggregates are embedded in an extracellular polymeric substance (EPS) that maintains a hydrated surface around the bacteria and thus helps them to survive under conditions of desiccation (Condemine et al., 1999). Motility and chemotaxis are essential for *D. dadantii* when searching for favorable sites to enter into the plant apoplast. Therefore, we evaluated the consequences of the *hfq* or *proQ* mutations on motility and biofilm formation. The ability of the *hfq* and *proQ* mutants to swim was analyzed using capillary and soft agar assays, respectively. Incubation time during the capillary assays was short, so that we were able to overlook the impact of the growth defect between the *hfq* mutant and the WT strain. Soft agar assays were performed to test the motility of the *proQ* mutant, since similar growth rates were obtained for the *proQ* mutant and the WT strain. A motility index was calculated for both experiments (Figure 4A). It was equal to the motility of the mutant strain (number of cells in the capillary or size of the halo) divided by that of the WT strain. Both *hfq* and *proQ* showed reduced motility compared to the WT (38% and −22%, respectively). This reduced cell motility is in agreement with the behavior of *hfq* mutants of most pathogenic bacteria (Chao and Vogel, 2010; Vogel and Luisi, 2011; Sobrero and Valverde, 2012; Wagner, 2013; Updegrove et al., 2016).

**Figure 4:**
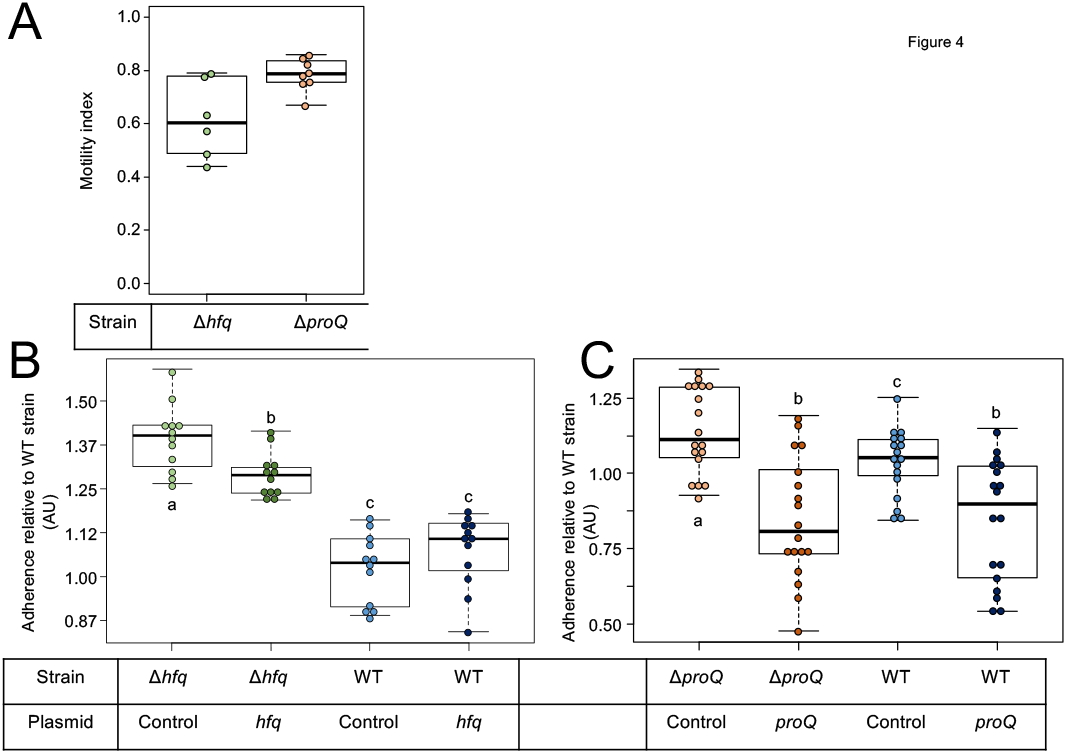
Impact of Hfq and ProQ on motility and biofilm formation. A, Motility indexes of the *hfq* and *proQ* mutants. Motility experiments were performed in capillary assays for the *hfq* mutant and in soft agar plates for the *proQ* mutant. The motility index was equal to the motility results of the mutant strain (number of cells in the capillary or halo size) divided by the results of the WT strain. B and C, impact of the *hfq* and *proQ* mutations on biofilm formation. Assays were carried out in multi-well plates. Data were normalized relative to the adherence of the wild-type strain. The effect of heterologous complementation is also showed. Quantification of the cells present in the aggregates and in the planktonic fractions for the different strains; a/b boxplot annotations highlight significant differences (*P* < 0.05, Wilcoxon Mann-Whitney test).

Flagellar motility negatively affects biofilm formation. Consequently, we monitored the consequences of the *hfq* or *proQ* mutations on the attachment of *D. dadantii* to the plastic coating of the microtiter plate well. From a metabolic point of view, biofilm formation reflects the trade-off between motility and exopolysaccharide (EPS) production. This trade-off was clearly unbalanced infavor of EPS production in the *hfq* mutant (*p*-value =7.4e-7) and less severely so in the *proQ* mutant (*p*-value =3.8e-2) (Figure 4B & C). Complementation experiments demonstrated that *hfq* expressed from an episome did not significantly reduce the increased biofilm forming capacity of the *hfq* mutant (Figure 4B). In contrast, expression of episomal *proQ* slightly decreased the biofilm formation capacity of the *proQ* mutant and WT strain (Figure 4C). These data suggest that the two RNA chaperones play different roles in *D. dadantii* biofilm formation.

### 3.6 Transcripts of late virulence factors and early virulence determinants are impaired in the *hfq* and *proQ* mutants

Hfq and ProQ act post-transcriptionally, so we evaluated the mRNA amounts of various virulence genes in the *hfq* and *proQ* mutants by qRT-PCR. For genes mostly involved in the early stage of infection, we selected *fliC* which encodes flagellin, *rhlA* whose product is involved in the synthesis of a biosurfactant for swarming motility, and *bcsA* which encodes a protein involved in the production of cellulose fibrils important for adherence. Concerning late virulence genes, we retained *pelD* and *pelE* that encode pectate lyases, *prtB* and *prtC* that encode metalloproteases, *celZ* that encodes a cellulase, *outC* that encodes a compound of the type II secretion system which secretes pectinases and cellulase, *kdgK* that encodes the KDG kinase involved in the catabolism of polygalacturonate, and *hrpN* that encodes harpin which elicits the hypersensitive response. In line with the observed phenotypes, the RNA amounts of most genes were reduced in both mutants, much more drastically so in the *hfq* mutant than in the *proQ* mutant (Figure 5). The greater adherence of the *hfq* mutant was also correlated with the higher *bcsA* RNA amounts compared to the WT. However, *celZ* RNA amounts were similar in the *hfq* mutant and in the WT. Therefore, the reduced cellulase activity was not dependent on the *celZ* RNA amount but it could be partially due to reduced cellulase secretion because the *outC* RNA amount was low in the *hfq* mutant, or to decreased CelZ translation. In the *proQ* mutant, the amount of *celZ* transcripts was reduced, but the amount of *outC* transcripts was not. Taken together, these results indicate that most of the observed phenotypes were correlated to a decrease in the mRNA amounts from virulence genes.

**Figure 5:**
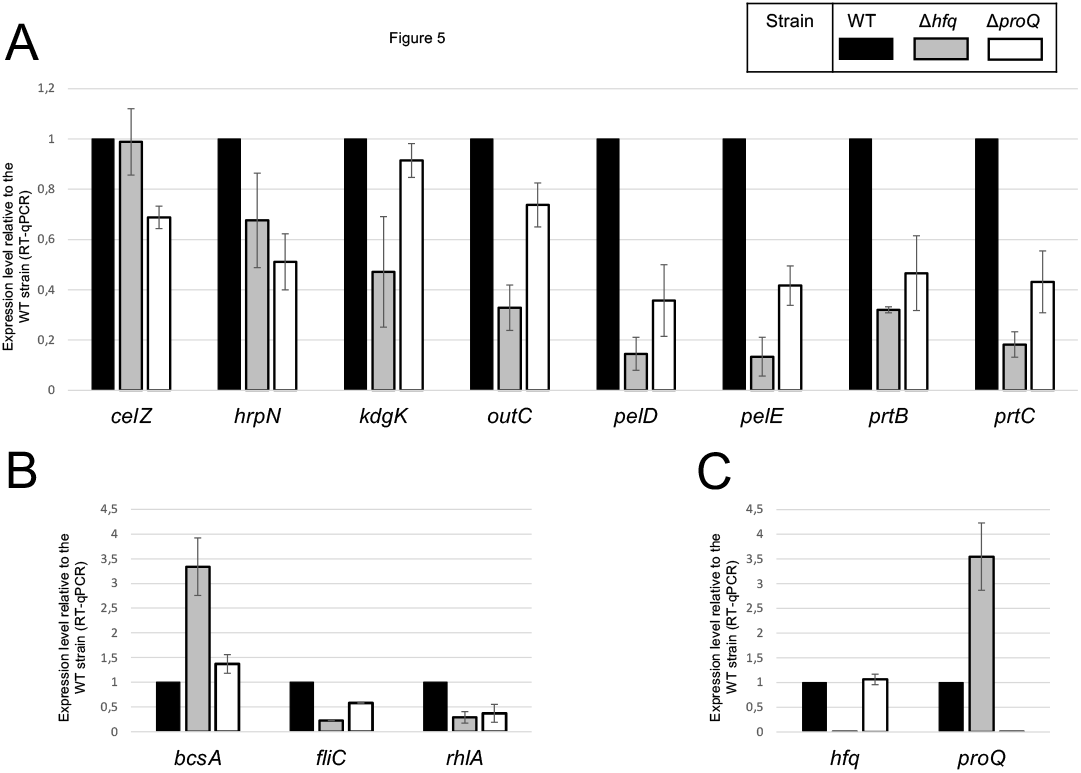
RNA amounts in the *hfq* and *proQ* mutant strains. Gene expression levels relative to the WT strain were evaluated in the two mutants by RT-qPCR. A, Genes encoding late virulence factors or associated with late virulence factors; B, Genes encoding early virulence factors such as adherence and motility factors; C, Expression levels of *hfq* and *proQ* were measured in the different mutants.

### 3.7 The effects of RNA chaperones on virulence partially overlap

The absence of Hfq or ProQ impaired virulence and modified the production of similar virulence factors. Consequently, we evaluated the behavior of a mutant inactivated for both *hfq* and *proQ* and assessed whether *hfq* and *proQ* could restore virulence in the double mutant. The virulence of the *hfq proQ* double mutant and mutant complemented by either *hfq* or *proQ* was tested on chicory leaves. The mutants were asymptomatic whatever the complementation 24 h post infection, except the mutant *proQ*, that caused reduced soft rot symptoms as noticed earlier (Figure 2). However, 48 h post infection, the *proQ* gene inserted in the *hfq proQ* mutant produced a weight of macerated tissues similar to the weight observed with the *hfq* mutant, whereas *hfq* complementation of the double mutant only slightly restored soft rot symptoms (Figure 6).

**Figure 6:**
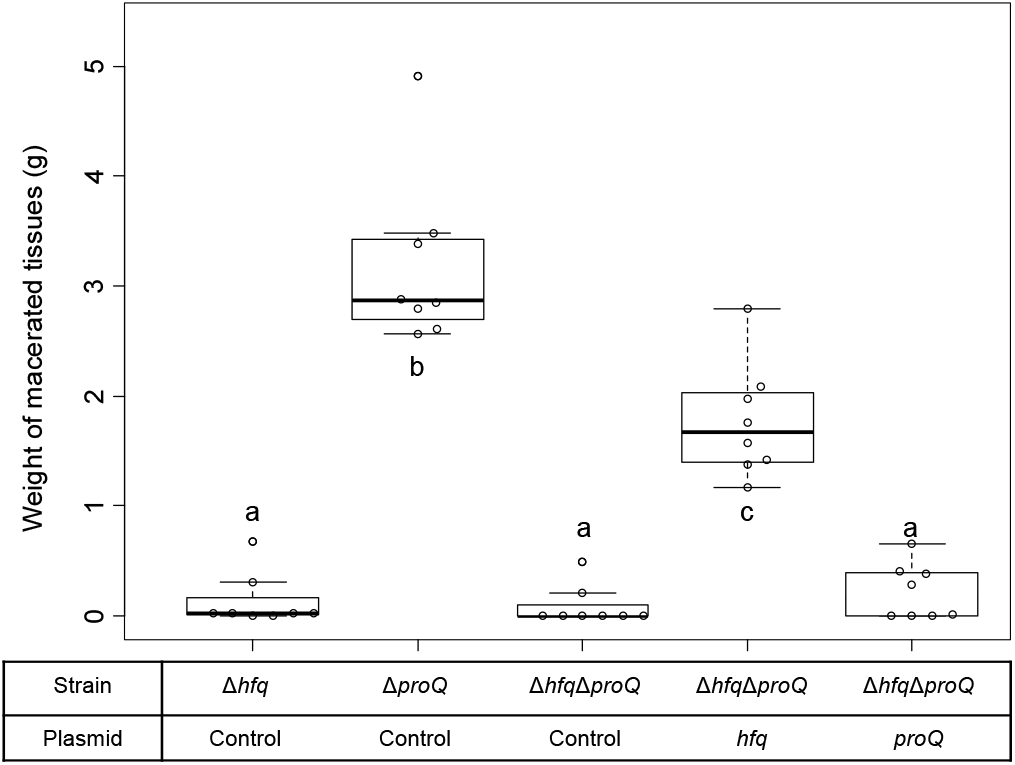
*D. dadantii* virulence assays 48h post infection. Virulence was evaluated on the single mutants, the double mutant, and the double mutant complemented by Hfq or ProQ. Chicory leaf assays were performed as described in the Materials and methods section with an incubation time of 48h, and the weights of macerated tissues were measured.

Late and early virulence factors were also impaired in the double mutant, as expected. Compared to the *hfq* mutant, the double mutant showed a similar, perhaps higher reduction of protease, cellulase and pectinase activities (Figure 7A). Motility was also reduced, and adherence was increased compared to the WT (Figure 8A and C). In line with these phenotypes, the expression levels of the corresponding genes were modified: *prtC, pelD* and *fliC* RNA amounts decreased by at least 5 folds in the double mutant, and *hrpN, celZ* and *outC* RNA amounts decreased by about 2 folds (Figures 7B, 8 and S4). Overall, the expression levels of *outC, pelD, bcsA* and *fliC* were similar in the double mutant and in the *hfq* mutant, whereas the expression levels of *prtC celZ* and *hrpN* decreased more than in the *hfq* mutant (Figures 7B, 8B and S4).

**Figure 7:**
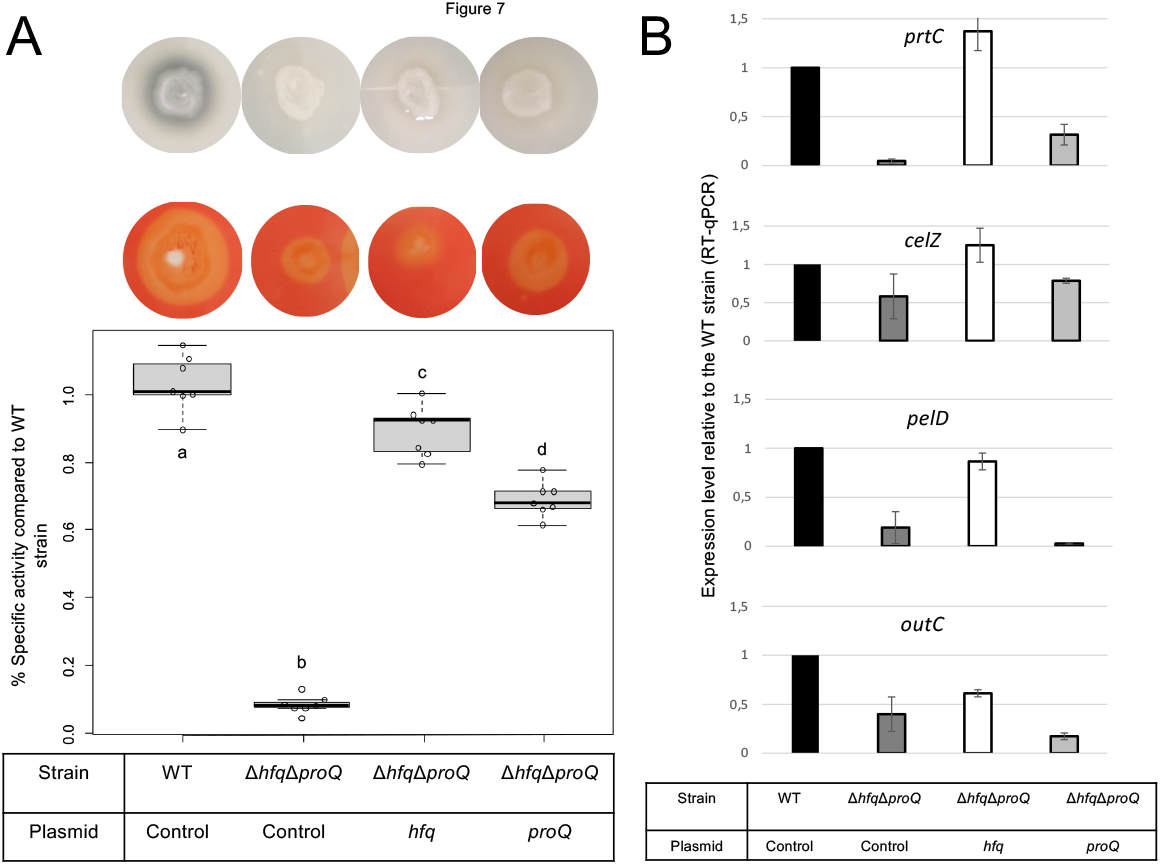
Impact of the double mutant on early virulence factors. Phenotypes and expression levels were analyzed in the double mutant strain and the double mutant strain complemented by Hfq or ProQ. a/b/c/d boxplot annotations highlight significant differences (*P* < 0.05, Wilcoxon Mann-Whitney test). A, Motility was evaluated on semi-solid LB agar plates; B and D, Expression levels of genes were measured by RT-qPCR and compared with the wild type; C, Adherence was evaluated and compared with the wild-type strain.

**Figure 8:**
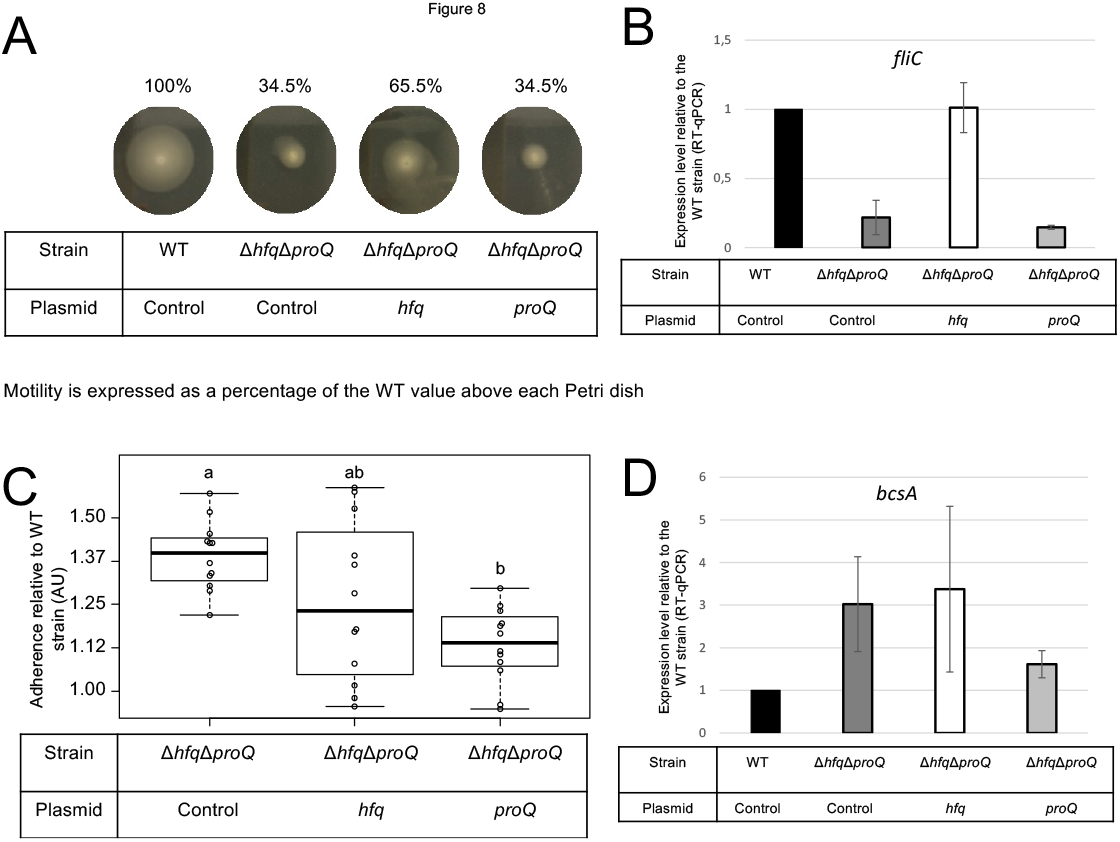
Impact of the double mutant on early virulence factors. Phenotypes and expression levels were analyzed in the double mutant strain and the double mutant strain complemented by Hfq or ProQ. A, Motility was evaluated on semi-solid LB agar plates; B and D, Expression levels of genes were measured by RT-qPCR and compared with the wild type; C, Adherence was evaluated and compared with the wild-type strain. a/b boxplot annotations highlight significant differences (*P* < 0.05, Wilcoxon Mann-Whitney test).

The protease, pectinase and motility phenotypes were restored by complementation with Hfq. Accordingly, *prtC, pelD* and *fliC* RNA amounts increased significantly in the double mutant strain complemented by Hfq (Figures 7B and 8B). However, the cellulase phenotype was not complemented by the addition of Hfq, even if the expression level of *celZ* was restored to a level similar to those measured in the WT or the *hfq* mutant (Figure 7). The adherence phenotype and the expression level of *bcsA* were not complemented by an *hfq* gene expressed from a plasmid (Figure 8). ProQ complementation restored protease, cellulase and pectinase activities and adherence, but not motility. In accordance, the expression level of *fliC* was similar to the levels in the double mutant (Figure 8B), but the expression levels of *bcsA, prtC, celZ* and *hrpN* increased in the complemented strain compared to the non-complemented double mutant (Figures 7, 8 and S4). Interestingly, the expression levels of genes *outC* and *pelD* were unexpectedly similar to those observed in the double mutant although phenotypes were restored.

We measured the expression levels of *hfq* and *proQ* in the different mutants. As expected, *hfq* and *proQ* expression was not detected in the respective mutant strains or in the double mutant, but a 3-fold increase was observed for the gene expressed from the plasmid. *proQ* expression level increased in the strains defective in Hfq production, *i.e*. 3.5 fold in the single *hfq* mutant and around 12 fold in thedouble mutant complemented by *proQ*. These results can be explained by an additive effect of the *hfq* mutation and of *proQ* overexpression from the plasmid (Figure 1 and S4).

Overall, virulence assays, phenotypes and expression level measurements suggest that the two RNA chaperones Hfq and ProQ exert their effects via partially overlapping but different sets of targets.

## 4 Discussion

We investigated the influence of the two RNA chaperones Hfq and ProQ on the virulence of the bacterial plant pathogen *D. dadantii*. Inactivation of the genes encoding these two chaperones led to lower production of cell-degrading enzymes acting as major virulence factors during *D. dadantii* pathogenic growth, and accordingly lower pathogenicity. Furthermore, both mutations altered osmotic stress tolerance and cell motility. However, the same mutations elicited different effects on cell growth and biofilm formation. Phenotypes were mostly correlated with altered expression of genes encoding virulence factors (*hrpN, outC, pelD, pelE prtC, prtB*), motility components (*fliC* and *rhlA*) and adherence components (*bcsA*), except *celZ* expression in the *hfq* mutant. Expression levels generally decreased less following inactivation of *proQ* than following inactivation of *hfq*, but both RNA chaperones affected similar virulence factors. So far, the involvement of ProQ in virulence has been only reported in *Salmonella* where it regulates motility directly by downregulating *fliC* mRNA and represses or activates the expression of virulence genes (genes localized in SPI and SPII, respectively). Accordingly, infection by a *Salmonella proQ* mutant resulted in a decreased invasion rate in eukaryotic cells (Westermann et al., 2019). The present study reports for the first time the involvement of ProQ in the virulence of a plant-pathogenic bacterium. In *D. dadantii*, the amount of *fliC* mRNA decreased in the *proQ* mutant, but major virulence genes (*pel, prt* and *cel*) were repressed by ProQ. Differences in *proQ* mutant behavior were also highlighted by comparing mutant strains in *E. coli* and *D. dadantii*: the *proQ* mutant displayed impaired biofilm formation in *E. coli*, whereas it displayed increased adherence in *D. dadantii* (Sheidy and Zielke, 2013). Overall, this illustrates the species specificities of the ProQ regulatory networks, as previously described (Attaiech et al., 2016; Holmqvist et al., 2018; Smirnov et al., 2016; Westermann et al., 2019). Specificities could be a consequence of a rather distinct sRNA landscape produced by these bacterial species, where only small numbers of sRNA homologs overlap (Leonard et al., 2019). In contrast to ProQ, Hfq proteins have already been reported to significantly reduce virulence in several plant-pathogenic bacteria like *A. tumefaciens, E. amylovora* and *P. carotovorum*. However, the role of Hfq still remains only partially understood (Wilms et al., 2012) Zeng et al., 2013; Wang et al., 2018). The phenotypes of the *D. dadantii hfq* mutant are similar to those of the *P. carotovorum hfq* mutant: *hfq*-defective strains present a decreased growth rate, low cellulase, protease and pectinase production, and altered biofilm formation and motility.

One interesting feature highlighted by this study is the interplay between the two RNA chaperones. The mitigated virulence of the double mutant was only slightly complemented by Hfq or ProQ, so we evaluated the ability of Hfq and ProQ to cross-complement each other regarding mitigation of virulence. ProQ partially complemented the *hfq* mutant, whereas episomal *hfq* did not complement the *proQ* mutant (Figure S5). Overall, these results indicate that these two RNA chaperones might exert their effects via partially overlapping but different sets of targets. Although first reports indicate that the RNAs bound by ProQ generally differ from those bound by Hfq (Holmqvist et al., 2018), recent studies have demonstrated an unexpected overlap of the sets of Hfq and ProQ targets in *Salmonella* and *E. coli*, with 30% of overlapping interactions (Westermann et al., 2019; Melamed et al., 2020). In line with these results, we identified potential ProQ-specific targets such as *celZ*, but also overlapping targets - the *fliC, bcsA, pel* and *prt* genes. Nonetheless, the expression of the target genes was more highly impacted in the double mutant than in each single mutant, indicating putative additive effects of the two proteins. Additionally, analyses of the expression levels of these target genes in the double mutant complemented by each protein highlighted 3 classes of genes: (i) *hrpN-* like genes, whose expression level is restored by Hfq or ProQ, (ii) genes whose expression levels are restored at least partially only by Hfq (e.g. *fliC, prtC, outC* and *pelE*), and (iii) genes whose expression levels are partially restored only by ProQ (e.g. *bcsA*). Although further studies aimed at identifying the direct targets of Hfq and ProQ in *D. dadantii* by *in vivo* crosslinking will clarify whether the virulence functions governed by ProQ represent a subset of those governed by Hfq, these data reinforce the assumption that the two proteins could have independent competing or complementary roles. ProQ and Hfq could be involved in different regulatory cascades, with branches converging at identical targets. Alternatively, both proteins could interact with a same mRNA. The site of interaction would not necessarily be identical: ProQ recognizes its targets in a sequence-independent manner, through RNA structural motifs found in sRNAs and internal to the coding sequence region or at the 3’ end of mRNAs (Holmqvist et al., 2018); whereas Hfq interacts with nascent transcripts in the 5’-UTR of the target RNA (Kambara et al., 2018). However, the two proteins could outcompete each other at a given terminator since they both have the propensity to bind intrinsic terminators of RNAs (Holmqvist et al., 2016).

Finally, it is noteworthy that an increased level of *proQ* was measured in the *hfq* mutant and in the double mutant complemented by ProQ. This is of importance because the impact of ProQ on growth and biofilm production is known to be highly dependent on the amount of ProQ protein produced in the cells. Taken together, these results indicate that in addition to the overlapping, complementary, competing or even additive roles of these two RNA chaperones, Hfq could indirectly or directly regulate ProQ production. Further analyses of the complex regulatory network of Hfq and ProQ should take this possible regulation into account.

## 7 Acknowledgments

S. Leonard received a doctoral grant from the French Ministère de l’Education nationale de l’Enseignement Supérieur, de la Recherche et de l’innovation. The authors thank Carlos Blanco for providing the *hfq* mutant, Pauline Héritier for helpful technical assistance and Georgi Muskhelishvili for helpful discussion. The authors also thank Annie Buchwalter for her careful correction of English language.

## 9 Supplementary Material

Table S1 to S4 in file named SupplementaryMaterials_Leonard.pdf

Figures S1 to S5 in file named SupplementaryMaterials_Leonard.pdf

