## Supplementary_Data for "RNA chaperones Hfq and ProQ play a key role in the virulence of the plant pathogenic bacterium *Dickeya dadantii*"

### Supplementary tables

TableS4: List of primers used in qPCR

| Gene names | Forward primers | Reverse primers |
|---|---|---|
| *bcsA* | CCCGATGGACAGTGAAAAAC | GGCGATAAACAACCCAATGC |
| *celZ* | TGCCGCTCTCTTATTTGGAT | CCCCAGCCATTATTACTCCA |
| *fliC* | CCCAGACCAACCTGAACAAA | TACCTTCAGCGGTCTGAACC |
| *hfq* | TAATGGCATCAAGCTGCAAG | TCAGCGTCATCACTTTCCTG |
| *hrpN* | TACGATTAAAGCGCACATCG | GTATTGAGCGACGCACCAAG |
| *kdgK* | AACACCGCCGTCTACATTTC | GGCATCGTTACGCCAGTAGT |
| *outC* | CTGCTGATGCTGCTCTTTTG | AGAAACGCCGAATAGCGTAA |
| *pelD* | TTGTGGAAGGTAACGCGCAGTTTG | ATGGCAAATTCACCAACGGCTCTC |
| *pelE* | AGCGAATTCAAAGCAGCACT | GGCGTTTCGATGTACAGGTT |
| *proQ* | TCTCCGTCATCCGAAAAATC | GGAAGCCAGTTGTACCCTGA |
| *prtB* | AAAGCGGCAAATCTGACCTA | TTTTGATTGGGGCTGACTTC |
| *prtC* | ATGACGCTCAACACGCATTA | AGCTGACCGACTGCAGAAAT |
| *rhlA* | GCATATTTCCGATCCTGCAC | CCCAGGAAATCGACAGGATA |

Table S2: Plasmids used in this study

| Plasmids | Description | Source |
|---|---|---|
| pKD3 | $Cm^R$ | Lab collection |
| pGEM-T | Cloning vector, $Amp^R$ | Promega |
| pGEM-T-Δ*proQ*- BglII | pGEM-T with the proQ coding region containing BglII site | |
| pGEM-T-proQ::Cm | pGEM-T with the proQ coding region containing chloramphenicol resistance cassette | This work |
| pBBR1-mcs4 | $Amp^R$ | (Kovach et al., 1995) |
| pBBR1-mcs4::proQ | pBBR1-mcs4 containing proQ ±500bp | This work |
| pBBR1-mcs4::hfq | pBBR1-mcs4 containing hfq ±500bp | This work |

Table S3: Primers used for genetic constructions.

| Primer | Sequence (5'-3') | Description |
|---|---|---|
| P1 | GTAGCGCGTTACTGTTTGAGCG | Forward primer located 500bp upstream *proQ* |
| P2 | GCTCATCCACGTTTTGCGGCCC | Reverse primer located 500 downstream *proQ* |
| P3 | GGAGATCTGAAATTTCCTGATTACAACGG G | Diverging with end of *proQ*; contains BglII site |
| P4+P3' | CCCGTTGTAATCAGGAAATTTCAGATCTA CGGAGGCCAACCTGGGCATGAAC | Diverging with start of *proQ* + reverse complement of P3; contains BglII site |
| P5 | GCTAGCGTAGCGCGTTACTGTTTGAGCG | Forward primer located 500bp upstream proQ; contains NheI site |
| P6 | AAGCTTGCTCATCCACGTTTTGCGGCCC | Reverse primer located 500 downstream *proQ*; contains HindIII site |
| P7 | GCTAGCGTGTTCATCAGTTTGCGATTGC | Forward primer located 500bp upstream hfq; contains NheI site |
| P8 | AAGCTTCACCAGACGCGTCGCCAGATGG | Forward primer located 500bp downstream *hfq*; contains HindIII site |

Table: List of primers used in qPCR

| Gene names | Forward primers | Reverse primers |
|---|---|---|
| *bcsA* | CCCGATGGACAGTGAAAAAC | GGCGATAAACAACCCAATGC |
| *celZ* | TGCCGCTCTCTTATTTGGAT | CCCCAGCCATTATTACTCCA |
| *fliC* | CCCAGACCAACCTGAACAAA | TACCTTCAGCGGTCTGAACC |
| *hfq* | TAATGGCATCAAGCTGCAAG | TCAGCGTCATCACTTTCCTG |
| *hrpN* | TACGATTAAAGCGCACATCG | GTATTGAGCGACGCACCAAG |
| *kdgK* | AACACCGCCGTCTACATTTC | GGCATCGTTACGCCAGTAGT |
| *outC* | CTGCTGATGCTGCTCTTTTG | AGAAACGCCGAATAGCGTAA |
| *pelD* | TTGTGGAAGGTAACGCGCAGTTTG | ATGGCAAATTCACCAACGGCTCTC |
| *pelE* | AGCGAATTCAAAGCAGCACT | GGCGTTTCGATGTACAGGTT |
| *proQ* | TCTCCGTCATCCGAAAAATC | GGAAGCCAGTTGTACCCTGA |
| *prtB* | AAAGCGGCAAATCTGACCTA | TTTTGATTGGGGCTGACTTC |
| *prtC* | ATGACGCTCAACACGCATTA | AGCTGACCGACTGCAGAAAT |
| *rhlA* | GCATATTTCCGATCCTGCAC | CCCAGGAAATCGACAGGATA |

### Supplementary Figures

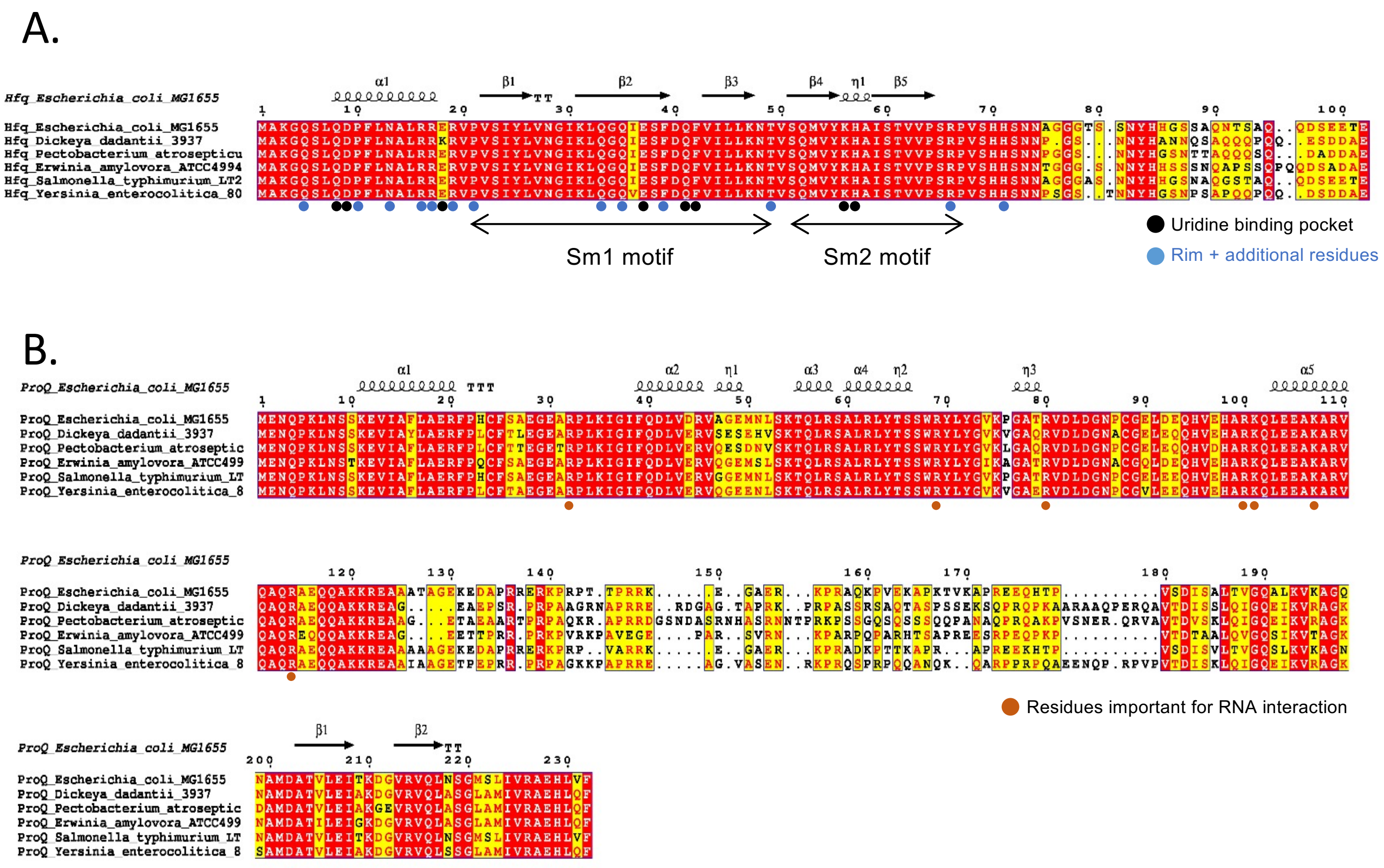

Figure S1: Sequence alignment and secondary structure prediction of Hfq (A) and ProQ (B) protein homologs identified in *Escherichia coli, Dickeya dadantii, Pectobacterium atrosepticum, Erwinia amylovora, Salmonella typhimurium* and *Yersinia enterocolitica.*

A.

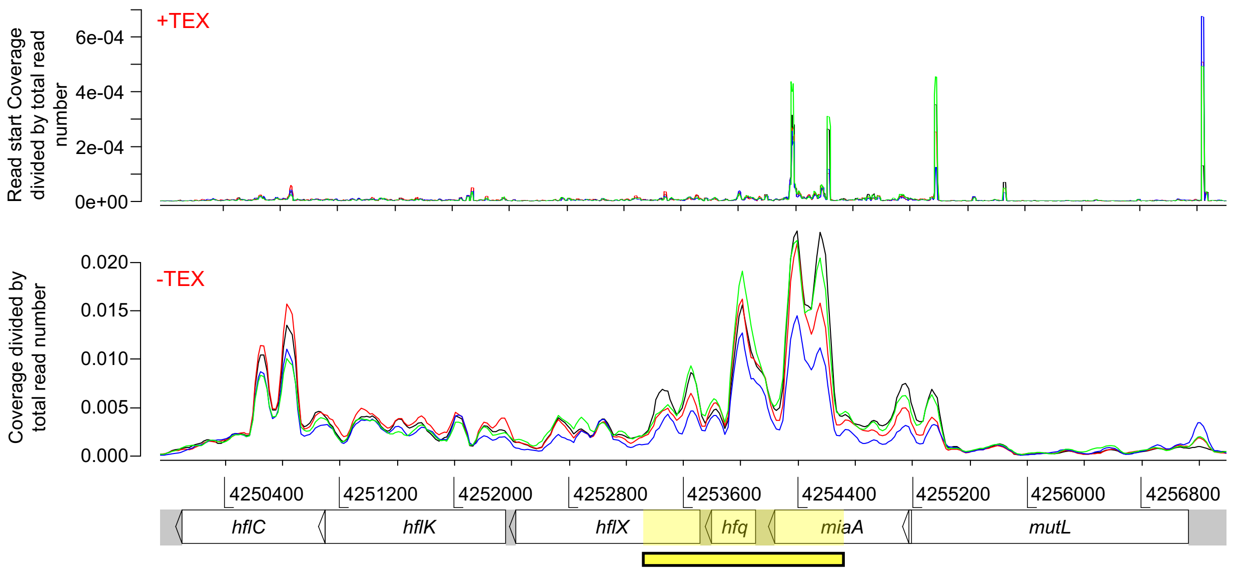

B.

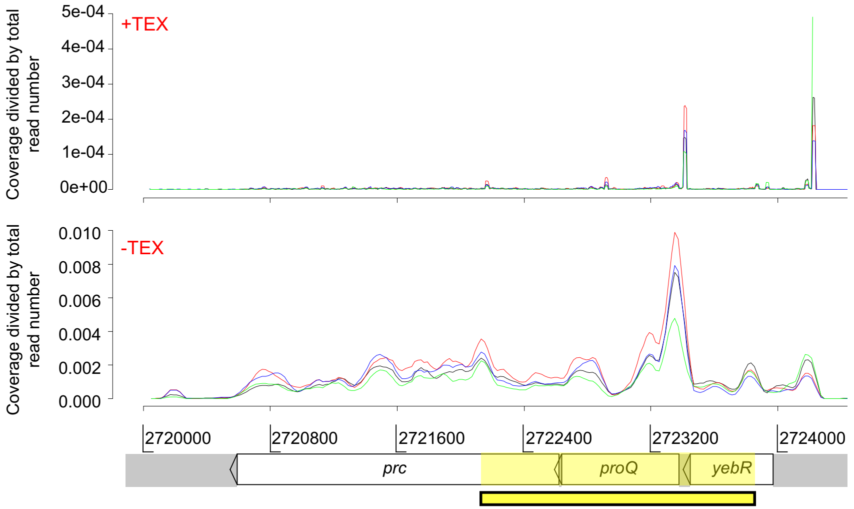

C.

| **Line color** | **Growth phase** | **Stress** |
|---|---|---|
| Blue | Stationary | No |
| Red | Exponential | No |
| Green | Stationary | Yes |
| Black | Exponential | Yes |

Figure S2: Genomic context and expression profiles of the *hfq* (A) and *proQ* (B) genes in *Dickeya dadantii* 3937. Genomic coordinates are given in the x-axis at the bottom of the figures. The normalized intensities (read coverage for the -TEX library and read start coverage for the +TEX library) are represented in the y-axis. Highlighted regions correspond to fragments used for plasmid complementation. Line colors represent the expression profiles, with sequencing conditions detailed in C.

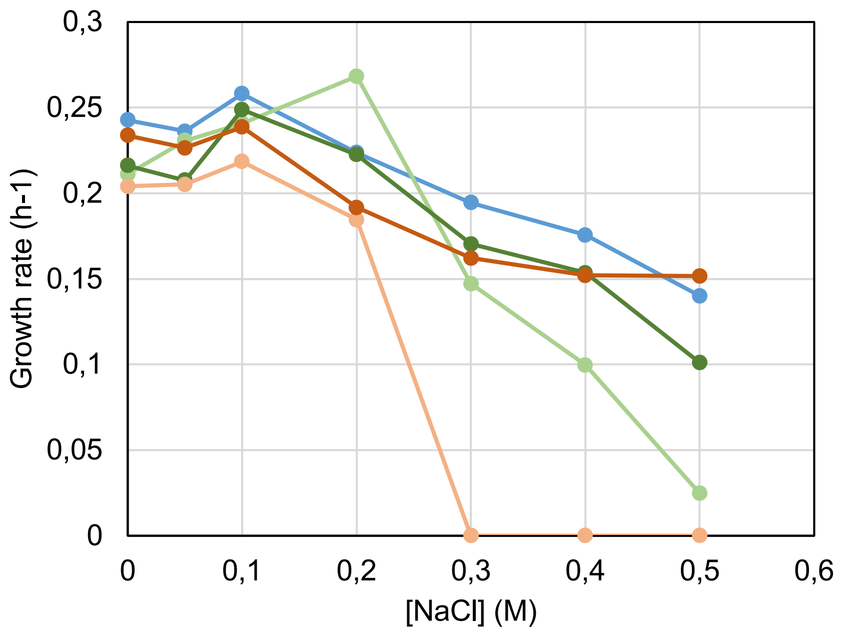

| | | | | | |
|---|---|---|---|---|---|
| Strain | Δ*hfq* | Δ*hfq* | WT | Δ*proQ* | Δ*proQ* |
| Plasmid | Control | *hfq* | Control | Control | *proQ* |

Figure S3: Growth of the wild type, mutant and complemented strains in M63 minimal medium with osmotic stress. Overnight bacterial precultures in M63 with sucrose as the carbon source and ampicillin (to maintained plasmid into the cells) were diluted to an $OD_{600}$ of 0.03 in a similar medium with $CaCl_2$ 0.1 mM + polygalacturonic acid (PGA) 0.025 % w/v. Osmotic stress was induced by adding different concentrations of NaCl in the medium. $OD_{600}$ measurements of the culture were made at regular intervals to determine growth rates.

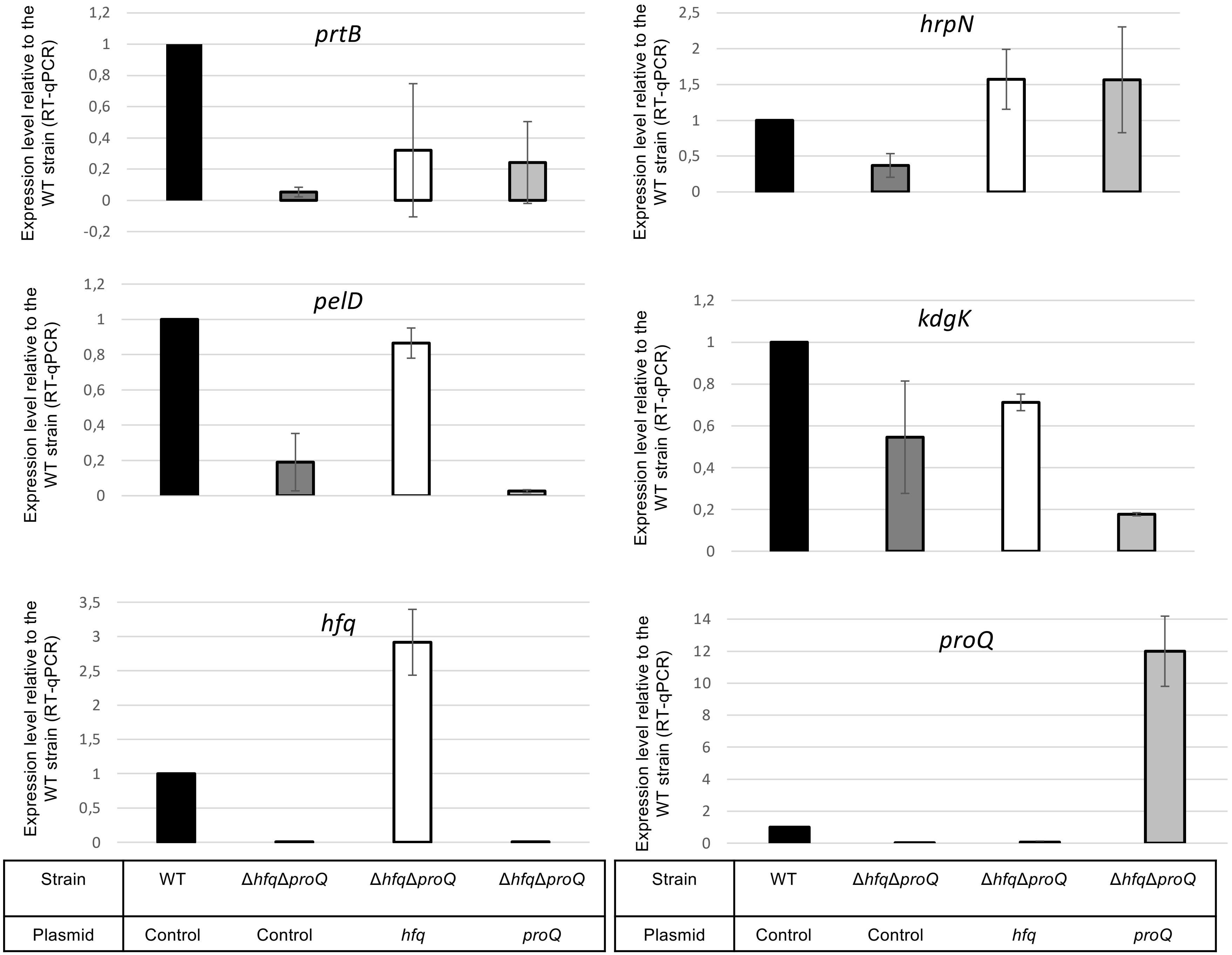

Figure S4: Expression levels in the double mutant strain and the double mutant strain complemented by Hfq or ProQ.

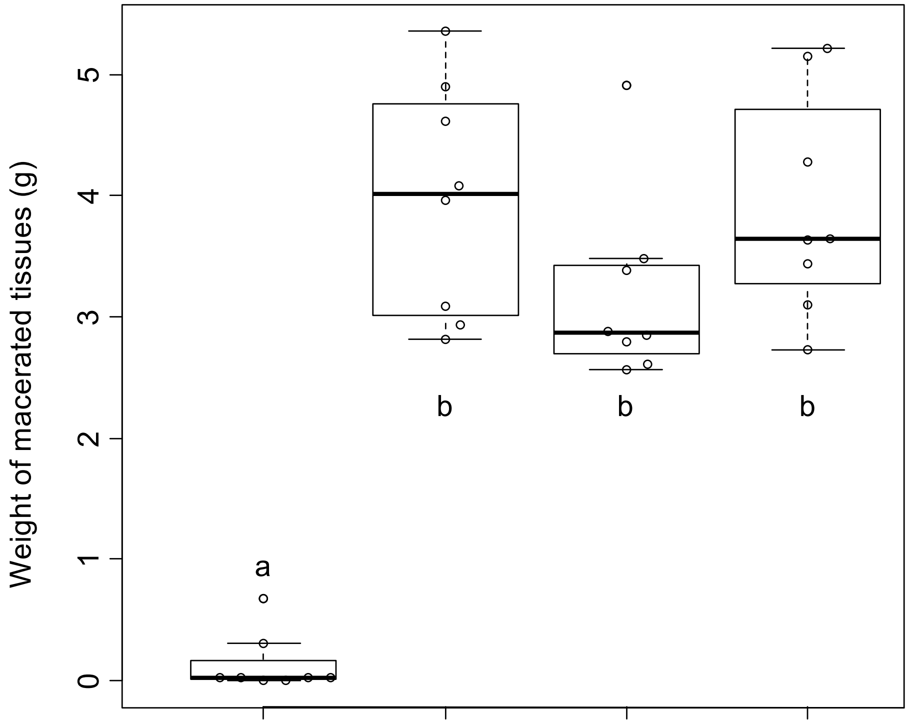

| Strain | Δ*hfq* | Δ*hfq* | Δ*proQ* | Δ*proQ* |
|---|---|---|---|---|
| Plasmid | Control | *proQ* | Control | *hfq* |

Figure S5: *D. dadantii* virulence assays 48h post infection. Virulence was evaluated on the *proQ* mutant with heterologous expression of *hfq,* and on the *hfq* mutant with heterologous expression of *proQ*. Chicory leaf assays were performed as described in the Materials and methods section with an incubation time of 48h, and weights of macerated tissues were measured.
